# Bacterial mobility and motility in porous media mimicked by microspheres

**DOI:** 10.1101/2022.11.08.515709

**Authors:** Diksha Shrestha, Jun Ou, Ariel Rogers, Amani Jereb, Yong Wang

**Affiliations:** Department of Physics, University of Arkansas, Fayetteville, Arkansas, USA, 72701; Cell and Molecular Graduate Program, University of Arkansas, Fayetteville, Arkansas, USA, 72701; Department of Environmental Resources Engineering, California State Polytechnic University Humboldt, Arcata, California, USA, 95521; Mechanical Engineering Program, California State Polytechnic University Humboldt, Arcata, California, USA, 95521; Materials Science and Engineering Program, University of Arkansas, Fayetteville, Arkansas, USA, 72701

**Keywords:** pore size, beads, geometric confinement, navigation

## Abstract

Bacterial motion in porous media are essential for their survival, proper functioning, and various applications. Here we investigated the motion of *Escherichia coli* bacteria in microsphere-mimicked porous media. We observed reduced bacterial velocity and enhanced directional changes of bacteria as the density of microspheres increased, while such changes happened mostly around the microspheres and due to the collisions with the microspheres. More importantly, we established and quantified the correlation between the bacterial trapping in porous media and the geometric confinement imposed by the microspheres. In addition, numerical simulations showed that the active Brownian motion model in the presence of microspheres resulted in bacterial motion that are consistent with the experimental observations. Our study suggested that it is important to distinguish the ability of bacteria to move easily – bacterial mobility – from the ability of bacteria to move independently – bacteria motility. Our results showed that bacterial motility remains similar in porous media, but bacterial mobility was significantly affected by the pore-scale confinement.

## Introduction

The motility and mobility of bacteria in heterogeneous porous media are important for pursuing nutrients and avoiding hazards in real-world settings [1–3]. For example, harmful bacteria navigate through porous structures in tissues to spread infections and invade human organs (Fig. 1A) [4, 5]. On the other hand, rhizosphere bacteria need to squeeze through pores in soil to, in many cases, beneficially impact the growth of plants (Fig. 1B) [6, 7]. Therefore, the motility and mobility of bacteria in porous media have attracted extensive attention in recent years, and a better understanding of the bacterial motility in porous structures is expected to shed light on modeling the spread of infections and developing useful strategies for biomedical and agricultural settings [1–3].

**Figure 1:**
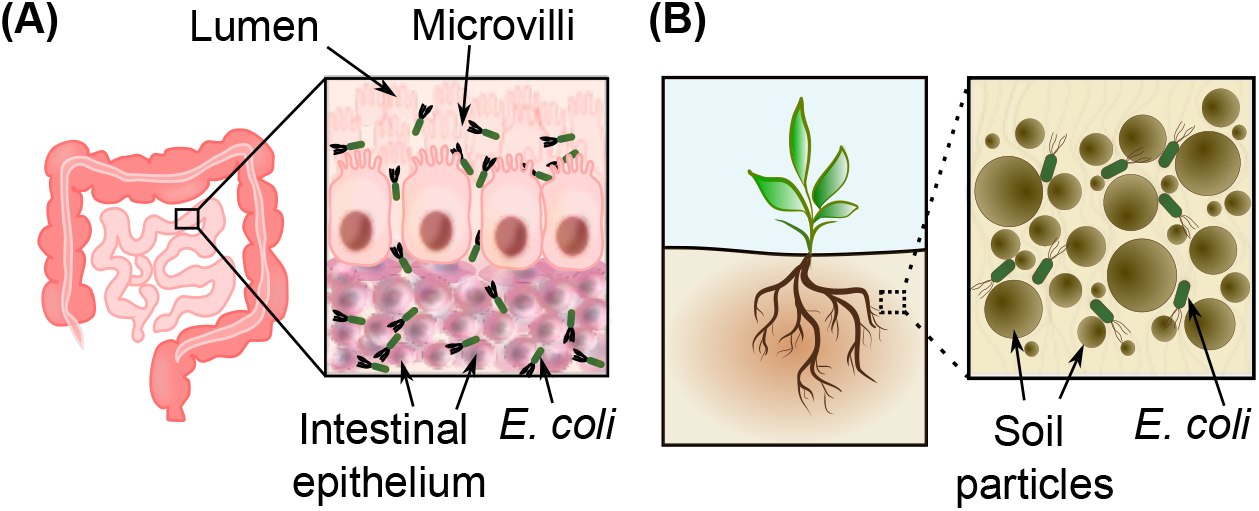
Examples of bacteria in porous media and structures: (A) *E. coli* bacteria in intestinal epithelium [4, 5] and (B) Rhizobacteria between soil particles in the rhizosphere [6, 7].

Previous theoretical, computational, and experimental studies have made exciting progresses towards understanding the motion and behavior of bacteria, and more generally, active Brownian particles and microswimmers, in porous media and/or complex, heterogeneous environments. For example, Zeitz et al. numerically investigated the dynamics of non-interacting active Brownian particles in a heterogenous environment with fixed and randomly distributed obstacles, and found that the long-distance transportation and trapping of the particles relied on the density of the obstacles [8]. Another computational study showed the avalanche dynamics of active Brownian particles in heterogeneous media [9]. In addition, phase separation was predicted from such systems from various numerical studies [10–13]. Various predictions from the theoretical and computational work have also been confirmed by experimental results. For example, Bhattacharjee et al. observed trapping and hopping and the avalanche dynamics for bacteria in 3D porous media made of micron-size hydrogel particles, which showed minimal light scattering and made the media transparent [2, 3]. On the other hand, the transparent hydrogel particles did not allow simultaneous study on how exactly bacteria interact with the pores. In addition, it remains unclear whether the motility of active swimmers was changed in porous media compared to in solution. Bhattacharjee et al. suggested that a fundamentally revised picture is required for the motility of active swimmers in porous media [2]; however, numerical studies implied that the unchanged underlying motility of active Brownian particles predicted the observed bacterial behaviors in heterogenous environments [8–10]. This controversy suggests that it may be necessary to distinguish between bacterial motility (how they move indepdently) and mobility (how they move easily), and to simultaneously and directly characterize both bacterial motion and pores in heterogeneous media.

In this study, we investigated the motion of *Escherichia coli* (*E. coli*) in microsphere-mimicked porous media. The rationale of using microspheres is two-fold. First, it is convenient to create a random, heterogeneous, pore-scale confining environment at different densities by controlling the concentration of microspheres – such a strategy has been used previously [14]. Second, the microspheres can be imaged and localized simultaneously with moving bacteria [15], allowing us to quantify both bacterial motion and their local pore-environment properties. Using microspheres of 6 *μ*m diameter at different densities, we tracked the quasi-2D motion of *E. coli* bacteria, quantified the instataneous velocities and directional changes of the bacteria, and examined the dependence of bacterial motion on the density of microspheres. We also determined the dependence of the average velocity and directional change of bacteria on the geometric confinement imposed by the pores formed by microspheres. More importantly, we examined the instaneous geometric confinement experienced by invididual bacteria along their trajectories, and established the correlation between bacterial hopping/trapping in porous media and the geometric confinement imposed by the microspheres. Lastly, we performed computaional simulations on bacterial motion in the presence of microspheres, and determined how well the active Brownian motion model applied to describe the bacterial motion in porous media.

## Materials and Methods

### Bacterial growth and sample preparation

Bacteria were grown and prepared following previous work [16**?** –19]. Briefly, an *E. coli* K12-derived strain, expressing histone-like nucleoid structuring proteins fused to mEos3.2 fluorescent proteins [16**?** –19], was used in the current study. For each experiment, a single colony was inoculated into 5 mL of Luria Broth (LB) medium, supplemented with kanamycin and chloramphenicol (at final concentrations of 50 *μ*g/mL and 34 *μ*g/mL, respectively) [16**?** –19]. The liquid culture was grown at 37°C overnight in a shaking incubator at 250 RPM, followed by dilution in fresh LB medium supplemented with kanamycin and chloramphenicol to reach OD_600_ = 0.005. The new culture (25 mL) was grown at 32°C in the shaking incubator until OD_600_≈0.2, followed by centrifugation (1000 g, 10 min, 21°C), removal of supernatant, and resuspension in 2.5 mL of fresh LB medium(i.e., ~10× concentration) [15]. Microspheres were pre-mixed by adding *X* = 0, 6, 10, 14 and 18 *μ*L of the original microsphere solutions to 18 – *X μ*L of water, followed by centrifugation at 1000 g for 10 min, removal of 5 *μ*L of supernatant, and addition of 5 *μ*L of 4× LB. Lastly, bacterial samples for imaging were prepared by adding 2 *μ*L of the concentrated bacteria to the 18 *μ*L of the premixed microspheres. Note that the use of 5 *μ*L of 4× LB ensured that the concentrations of LB medium remained at 1× for all the samples (20 *μ*L for each). The samples were labeled and referred to as BAC (*X* = 0) or MS*X* (*X* = 6, 10, 14, and 18).

Prepared bacterial samples were loaded to imaging chambers for microscopic measurements (Fig. 2A–2C). The imaging chambers were constructed by microscope slides and coverslips [20]. Briefly, the slides and coverslips were cleaned in a sonicator sequentially with soap, 1 M sodium hydroxide, and 100% ethanol, with thorough rinsing of deionized water (>17.5 MΩ) between different cleaning and sonication steps [16, 17, 21**?**]. The cleaned coverslips and slides were glued together by strips of double-sided tapes, from which imaging chambers were formed (Fig. 2A) [20]. The chambers were coated with bovine serum albumin (BSA, 10 mg/ml, 30 min) and then rinsed by phosphate-buffered saline (PBS), followed by loading 20 *μ*L of the samples [20].

**Figure 2:**
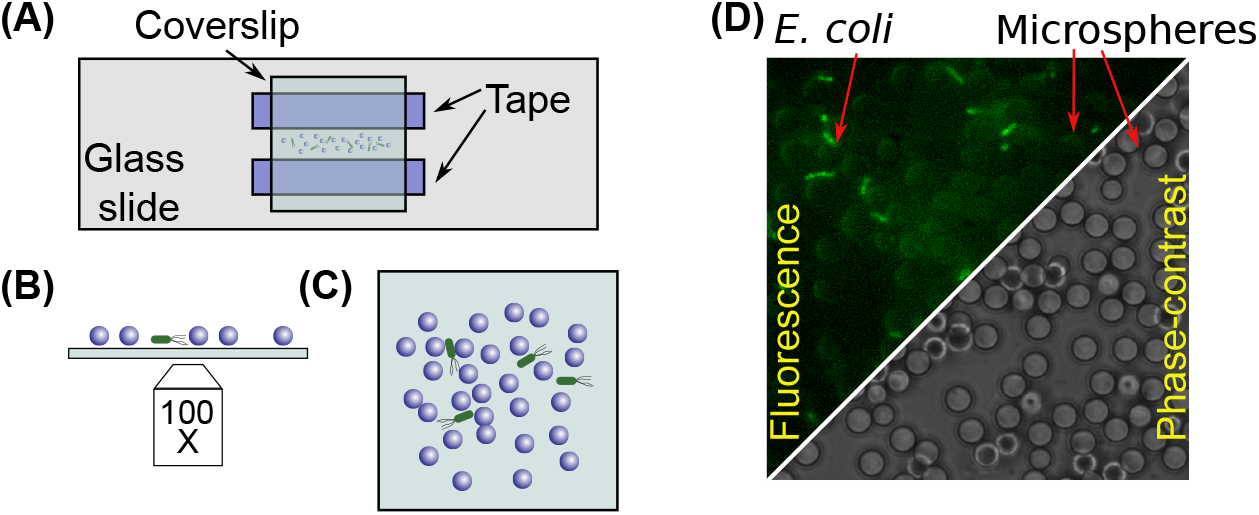
Illustration of sample preparation and imaging. (**A**) Schematic diagram of sample imaging chambers. (B) Sketch for imaging the bacteria and polystyrene microspheres: an 100× objective lens of an inverted microscope is focused at the proximity of the top surface of the coverslip. (**C**) Sketch of top view. (**D**) Examples of raw images of the bacteria in the presence of polystyrene microspheres, acquired with fluorescence microscopy (top-left) and phase-contrast microscopy (bottom-right).

### Phase-contrast and fluorescence microscopy

Phase-contrast microscopy and fluorescence microscopy were used to image the microspheres and bacteria (Fig. 2D) on an Olympus IX-73 inverted microscope with a 100× phase-contrast/fluorescence objective (NA=1.25, oil immersion). The microscope and data acquisition were controlled by *μ*manager [22, 23], while images were captured by an EMCCD camera (Andor Technology). With the 100× objective, the pixel size of of acquired images was 160 nm. For fluorescence imaging of the bacteria, the samples were first illuminated by handheld UV light (395 nm, ~20 mW) for 10 min, which activated the orange channel of mEos3.2 fluorescent proteins [24]. The bacteria were then imaged with the fluorescence microscope with laser-excitation at 532 nm (4 mW, iChrome MLE). The exposure time for the data acquisition was set to 15 ms, resulting in an actual interval of 39.6 ms between adjacent frames. The motion of bacteria were recorded as movies, and each movie consisted of 6,000 frames.

### Image processing and data analysis

Image processing and detection of bacteria and their trajectories were performed similarly to our previous work [15, 25]. Briefly, the acquired movies were first rescaled to half using ImageJ (i.e., from 512×512 to 256×256), resulting in an effective pixel size of 0.32 *μ*m [26, 27]. Next, for each frame of the rescaled movies, the background was removed using a rolling-ball algorithm with a ball size of 3 pixels, followed by smoothing twice using a Gaussian filter with a standard deviation of 1 pixel [15, 25, 28]. The background in the smoothed image was removed once again, followed by applying a threshold to obtain a black/white (BW) image [15, 25]. Edges were detected from the BW image using the Sobel filter, followed by dilating the edges by 1 pixel to fill possible gaps in the edges. Small objects with areas <25 pixels were removed, before performing a flood fill [15, 25, 29, 30]. The filled objects were eroded with 2 pixels, followed by removing small objects (area <25 pixels) [15, 25]. The resulting BW image was segmented into individual ones, which corresponded to the identified bacteria [15, 25]. The locations (*x*, *y*) of the bacteria, as well as their corresponding frames, were recorded and then linked into trajectories with *trackpy* [31], using a maximum displacement between adjacent frames of 5 pixels (1.6 *μ*m) and a memory of 3 frames. The trajectories of the bacteria 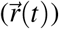 were saved and used for computations of the instantaneous velocities, 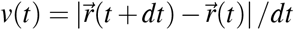, and directional changes, 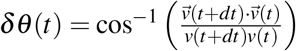, where *dt* = 39.6 ms is the time interval between adjacent frames [15, 32].

To locate the immobile microspheres, we first calculated the average image of each movie [15]. Note that moving bacteria disappeared while the immobile microspheres appeared in the averaged images, which facilitated the automated detection and manual label of microspheres. Although automated detection based on the circle hough transform using OpenCV [33] were successful for >95% of the microspheres, manual labeling of all the microspheres were performed in ImageJ to ensure 100% accuracy for the analysis in this study.

### Simulation of bacterial motion in the absence and presence of microspheres

The motion of bacteria was simulated using Python following the active Brownian motion model [15, 34]; however, the model was modified to account for the finite size of the bacteria. Briefly, *N_MS_* (0, 25, 50, 75, or 100) microspheres with a radius of *R_MS_* = 3 *μ*m were randomly placed in a region of 80×80 *μm*^2^ without overlapping with each other. The size of the simulation region (L = 80 *μ*m) was similar to the size of the field of view in our experiments (81.92 *μ*m). For simplicity and convenience, periodic boundary conditions were applied when a bacterium moved out of the region [15].

In the simulations, spherical bacteria (radius *R* = 0.5 *μ*m) were randomly placed in the region but outside the microspheres. Each bacterium was described by its location (*x*, *y*) and orientation *θ* [15, 34]. The position (*x_i_*, *y_i_*) and orientation (*θ_i_*) of a bacterium at time step *i* was tentatively calculated by its previous step, 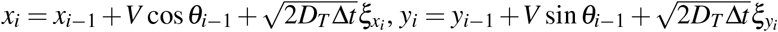, and 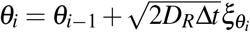, where *V* = 10 *μ*m/s is the deterministic linear velocity of the bacterium in the active Brownian model, Δ*t* = 0.02 s is the time step size, *D_T_* = *k_B_T*/6*πηR* is the translational diffusion coefficient, *D_R_* = *k_B_T*/8*πηR*^3^ is the rotational diffusion coefficient, *k_B_* is the Boltzmann constant, *T* = 300 K is the temperature, *η* = 10^−3^ Pa·s is the viscosity of the medium, and *ξ*’s are random numbers from a Gaussian distribution with a mean of 0 and a standard deviation of 1 [15, 34]. If the bacterium did not collide with any microspheres, the tentative values were taken. However, if the bacterium collided with a microsphere, the tentative bacterial location (*x_i_*, *y_i_*) was reflected at the effective boundary of the microspheres [15, 34], which was the circle centered at the center of the microsphere but with a radius of *R_MS_* + *R* = 3.5 *μ*m. For each simulation, 10,000 steps were performed, and the locations of the bacteria at each time step were recorded. The bacterial locations were then used for trajectory identification and quantitative analysis, as described for the experimental data.

## Results

### Bacterial trajectories

Several examples of bacterial trajectories are shown in Fig. 3, where each colored curve represents a trajectory of a single bacterium. Compared to the control (i.e., in the absence of microspheres; Fig. 3A), bacteria showed a higher frequency of large directional changes and confined motions (indicated by white arrows in Fig. 3B) in the presence of microspheres. We note that the large directional changes and confined motions of bacteria usually happened at the proximity of microspheres (Fig. 3).

**Figure 3:**
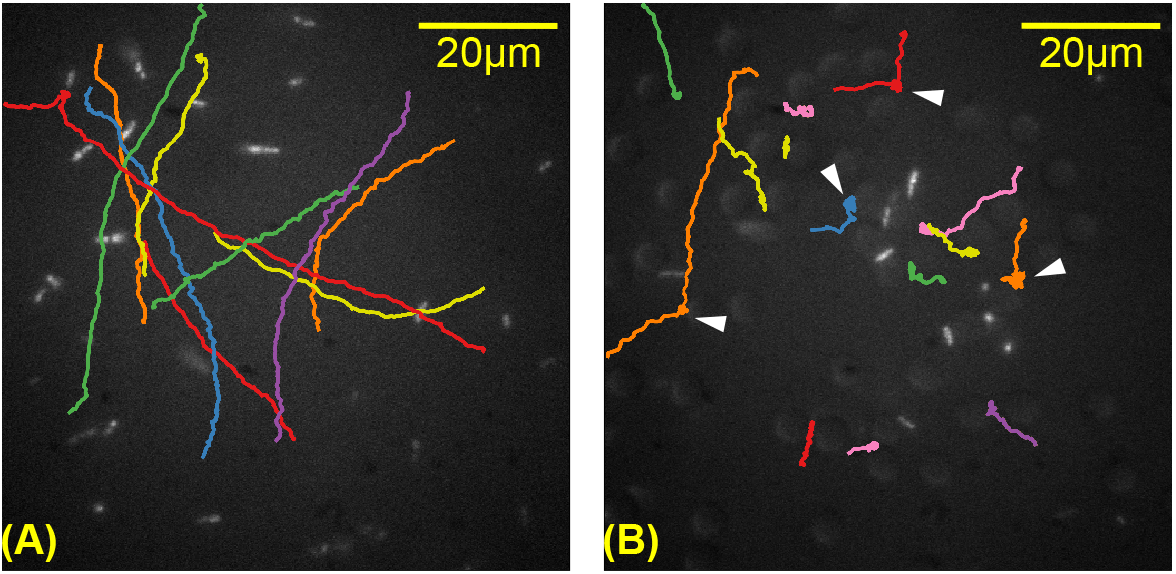
Examples of bacterial trajectories in the (**A**) absence and (**B**) presence of microspheres.

### Reduced velocity of bacteria due to the presence of microspheres

To systematically investigate the changes of of bacterial motion in microsphere-mimicked porous media, we varied the density of microspheres while keeping the concentration of bacteria constant. From the bacterial trajectories (Fig. 3), their instantaneous velocities were calculated. We then compared the average velocities of bacteria among the samples with different microsphere densities. As shown in Fig. 4A, we observed that the average velocities of bacteria in the presence of microspheres (MS6–MS18 in Fig. 4A) were significantly lower than that in the control (BAC in Fig. 4A). For example, the average velocity of bacteria in MS18 (3.9±0.7 *μ*m/s) was 60% lower than that in the control (9.9±1.1 *μ*m/s). It is worthwhile to note that the media among different samples were kept the same by (a) adding concentrated LB medium to all the samples, and (b) using a small volume of bacteria (2 *μ*L out of 20 *μ*L), suggesting that the observed difference in the bacterial velocity is due to the presence of microspheres. We also examined the distributions of the instantaneous velocities of bacteria in the absence and presence of microspheres (Fig. 4B). While the bacteria in the absence of microspheres showed a peak velocity around 10 *μ*m/s, the presence of microspheres shifted the velocity distributions to the left dramatically (Fig. 4B). A similar result was reported previously [2, 3].

**Figure 4:**
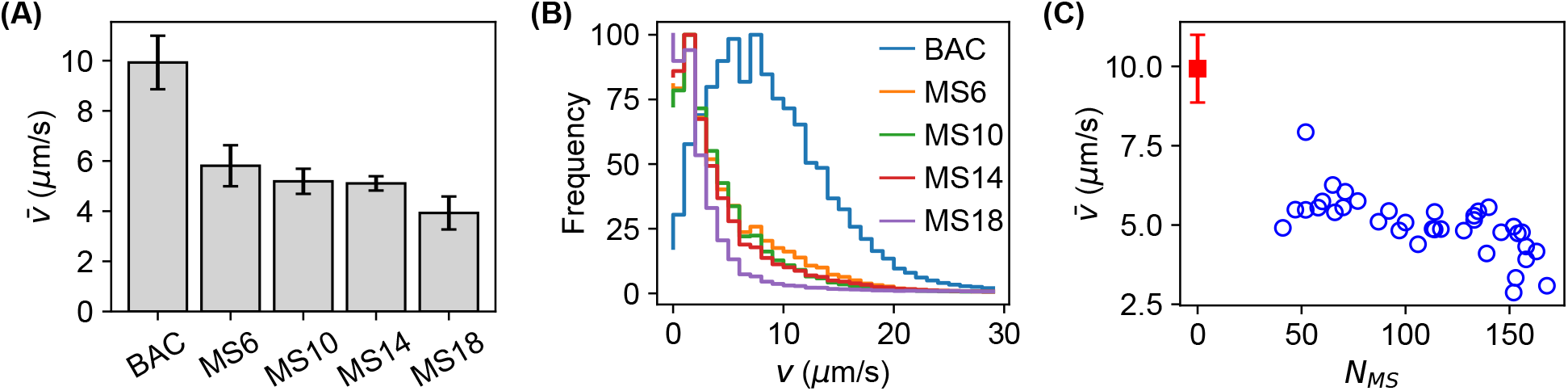
Bacterial velocity in porous media mimicked by microspheres. (**A**) Average bacterial velocity in samples of microspheres at different densities. (**B**) Distributions of bacterial instantaneous velocities of bacteria in different samples. (**C**) Average bacteria velocity of bacteria *vs*. the number of microspheres (*N_MS_*), where each blue circle represents data from a single movie (field of view).

As the microspheres were randomly distributed on the coverslip of the imaging chamber, the exact number of microspheres may be different in different fields of views of the same sample. This stochastic nature allowed us to more carefully examine the dependence of bacterial velocity on the density of microspheres. For each field of view, we counted the number of microspheres (*N_MS_*) and estimated the average velocity of bacteria in that field of view. A clear negative dependence was observed from this analysis: the average velocity of bacteria reduced as the number (and thus the density) of microspheres increased (Fig. 4C).

### Enhanced directional change of bacteria due to the presence of microspheres

The directional change of bacteria have been commonly used when studying bacterial motion. Similar to previous work, we estimated the change of bacterial direction per step/frame (*δθ*, Fig. 5A) from the bacterial trajectories in the absence and presence of microspheres at different densities. We observed that the average directional change of bacteria 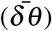 increased as the density of microspheres increased (Fig. 5B), supporting the qualitative observations from the trajectories (Fig. 3B). For example, the average directional change of bacteria in the MS18 sample (1.80±0.13 rad/frame) became 134% higher than that in the control without microspheres (0.77±0.04 rad/frame). A detailed look at the distributions of bacterial directional changes showed that the presence of microspheres caused that the bacteria took larger directional changes (Fig. 5C). The distribution of directional changes of bacteria was peaked around zero in the absence of microspheres (Fig. 5C), while it was maxed out around *π* for the bacteria in the MS18 sample. A similar result was reported previously [2, 3]. Furthermore, when counting the actual number of microspheres in each field of view, we observed a clear positive correlation: the average directional change of bacteria increased as the number (and thus the density) of microspheres increased (Fig. 5D).

**Figure 5:**
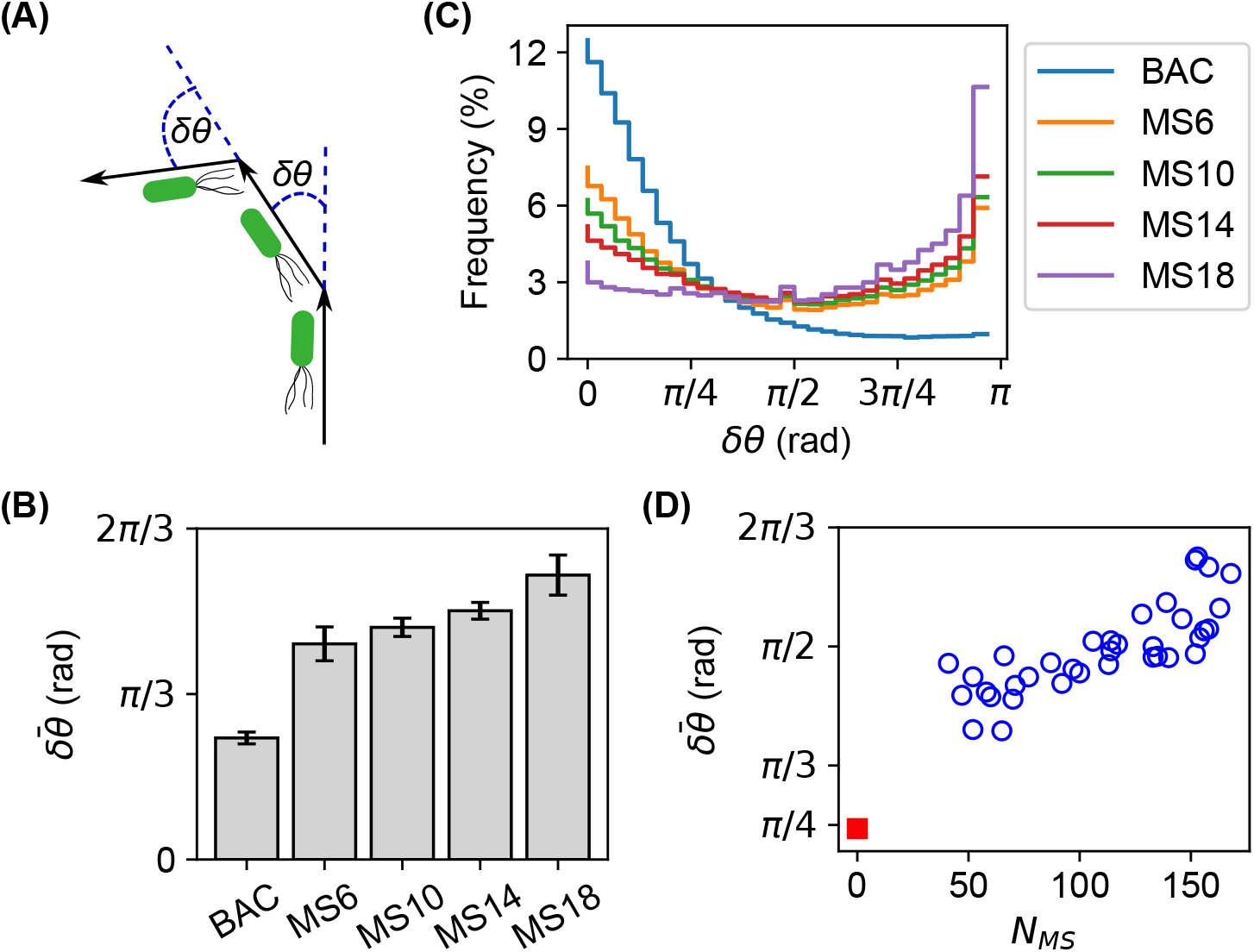
Directional change of bacteria in porous media mimicked by microspheres. (**A**) Change of bacterial direction 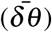 along the trajectory. (**B**) Average directional change of bacteria in samples of microspheres at different densities. (**C**) Distributions of directional change of bacteria in different samples. (**D**) Average directional change of bacteria *vs*. the number of microspheres (*N_MS_*), where each blue circle represents data from a single movie (field of view).

### Dependence of bacterial motion on geometric confinement

A significant advantage of using microspheres to mimic porous media is the capability of simultaneously localizing both the bacteria and the microspheres, which allowed us to determine the distance of the bacteria to the microspheres (*d_i_*) and the corresponding instantaneous velocity and directional change of the bacteria. As examples, we examined three geometric confinements: the surface of single microspheres, the gap formed between two microspheres, and the pore formed by three microspheres. These geometric confinements were mathematically characterized by 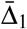 (the average of shortest distance, ⟨*d*_1_⟩), 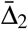 (the average of the shortest two distances, ⟨(*d*_1_ + *d*_2_)/2⟩), and 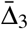 (the average of the shortest three distances, ⟨(*d*_1_ + *d*_2_ + *d*_3_)/3⟩), respectively, as shown in Fig. 6A. We observed that, below certain thresholds, the average velocity of bacteria increased linearly as 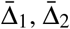 and 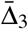 increased (Fig. 6B, D, and F). This observation suggested that stronger geometric confinements slowed down the bacteria. Above the thresholds, the average velocity of the bacteria reached plateaus, which corresponded to the average velocity of bacteria in the control without microspheres and geometric confinements (green regions in Fig. 6B, D, and F). Interestingly, we observed the thresholds of 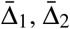, and 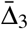 for reaching the plateaus were 3 *μ*m, 6 *μ*m, and 9 *μ*m, respectively.

**Figure 6:**
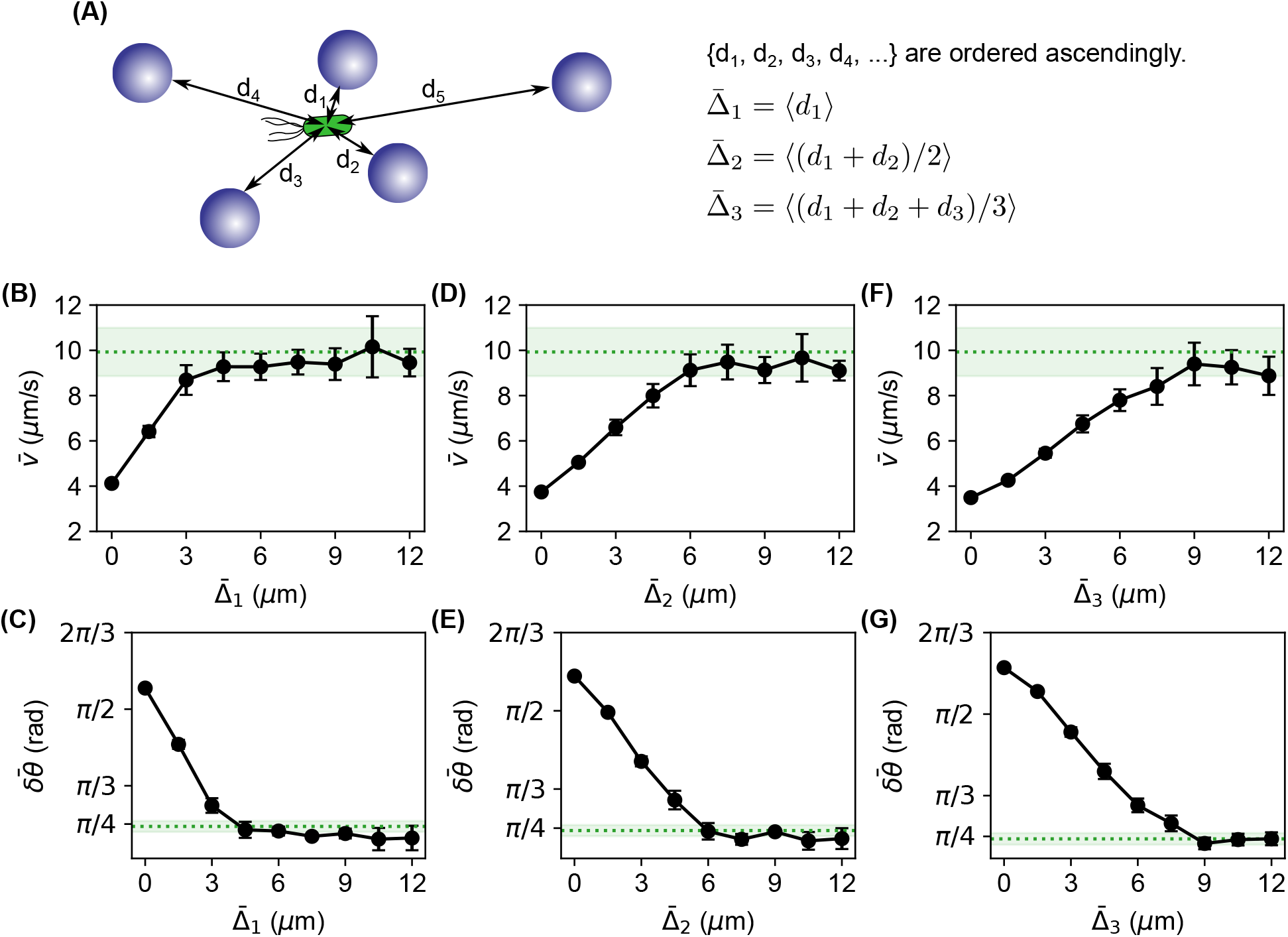
Dependence of the average velocity (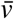) and the average directional change 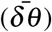 of bacteria on the geometric confinements. (**A**) Illustration of the geometric confinements to the bacteria due to the microspheres: the distance to the surface of single microspheres 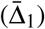, the gap formed between two microspheres 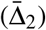, and the pore formed by three microspheres 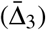. The confinements are estimated from the distance of the bacteria to the closest microspheres, with *d*_1_ to *d*_5_ numbered in ascending order. (**B, D, F**) Dependence of the avearge bacterial velocity 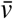 on the geometric confinements due to the microspheres. (**C, E, G**) Dependence of the avearge directional change of bacteria 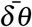 on the geometric confinements due to the microspheres.

In addition, we examined the dependence of the average directional change 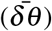 of bacteria on the geometric confinements. We observed that 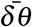 increased as the geometric confinements became stronger (i.e., lower 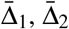, and 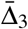, Fig. 6C, E, and G), suggesting that the bacteria tumbled more frequently and vigorously due to the geometric confinements. Furthermore, we found that, above the same thresholds from the average velocities (i.e., 3 *μ*m, 6 *μ*m, and 9 *μ*m for 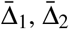, and 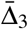, respectively), the average directional change of bacteria reach plateaus again, corresponding to the value of bacteria in the absence of microspheres and geometric confinements.

### Correlation between bacterial trapping and geometric confinement

It was reported that bacteria displayed hopping and trapping in porous media [2]; however, it is not clear how bacterial hopping/trapping correlates with the geometric confinement due to the porous media. Two examples of individual bacterial trajectories, velocities, directional changes, and the corresponding geometric confinements are shown in Fig. 7. We observed bacterial hopping and trapping from our microsphere-mimicked porous media (Fig. 7A and D), consistent with previous reports [2]. The bacterial velocity (*v*) reduced significantly when the bacteria were in the “trapping” mode, mostly below half of the average velocity of bacteria in the control without microspheres (Fig. 7A and D). Associated with the reduced velocity was the enhanced directional changes (*δθ*) and the larger fluctuations of *δθ* in the “trapping” mode. The larger fluctuations suggested that the bacteria may make efforts to overcome the “trapping” mode. More importantly, we found that reduced velocity of bacteria in the “trapping” mode correlated positively with the geometric confinement due to the microspheres. For example, the distance between the bacteria and the microspheres (e.g., 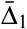 and 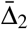) were much lower when the bacterial velocity reduced and the fluctuations of the directional change increased (Fig. 7A and D). Such geometric confinements were also obvious from the trajectories overlapped with the locations of the microspheres (Fig. 7B, C, E, and F). To statistically investigate the correlation between bacterial trapping and geometric confinement, we evaluated the Pearson’s correlation coefficients between the instantaneous velocity of bacteria and the corresponding gaps formed between two microspheres, and found that 80% of the trajectories showed positive correlations.

**Figure 7:**
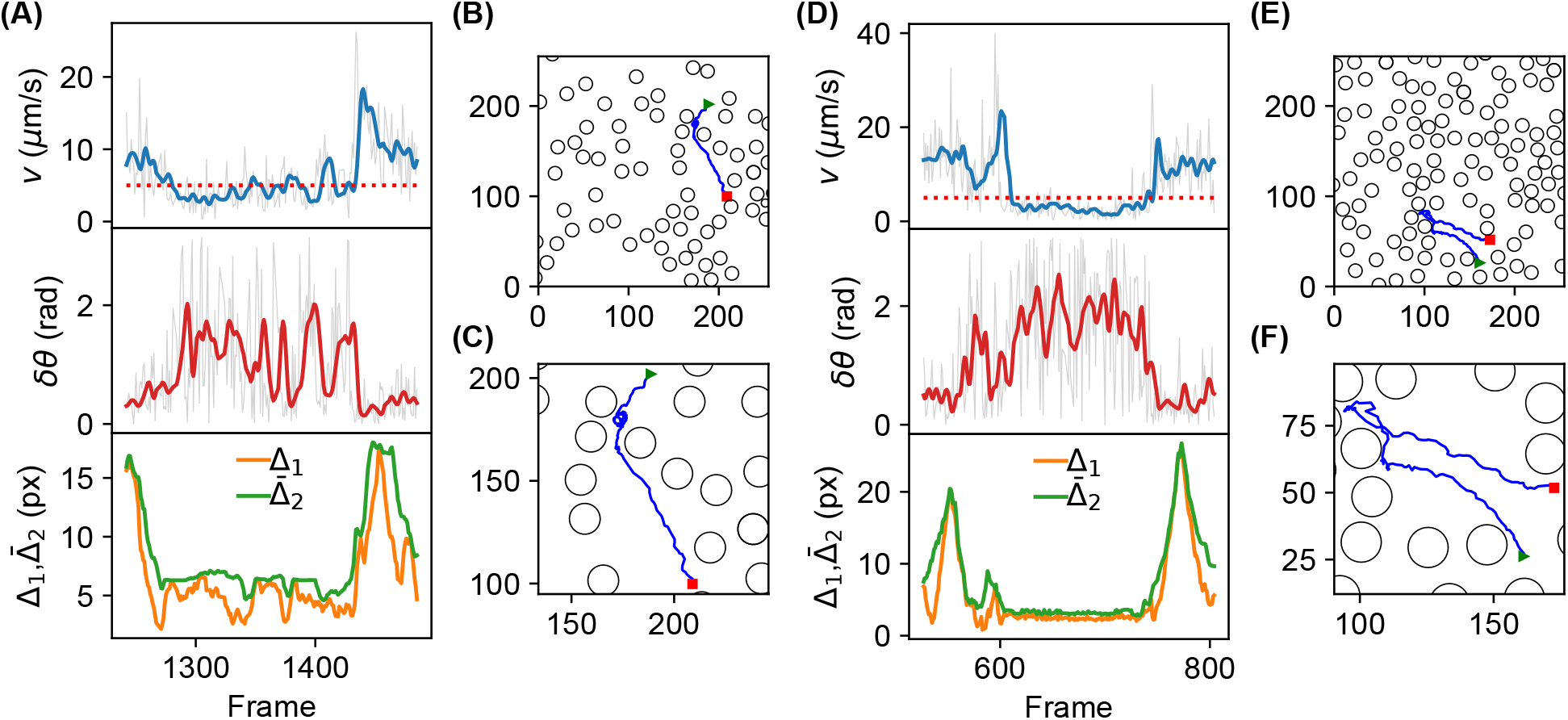
Examples of two individual trajectories of bacteria showing two phases: hopping and trapping. (**A, D**) The instantaneous velocity (*v*) and swimming direction (*δθ*) of a bacterium, and the gap sizes (represented by 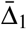 and 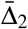), as functions of frame number (time). (**B, E**) The corresponding trajectory (blue line) of the bacterium in the presence of microspheres (black circles). The starting and ending points of the trajectory are represented by the green triangle and red square, respectively. (**C, F**) Zoom-in of the same data in (**B**) and (**E**), respectively.

### Simulation of bacterial motion in the presence of microspheres

To understand the observed motion of bacteria in the microsphere-mimicked porous media, we ran numerical simulations of bacterial motion in the absence or presence of microspheres at different densities. The bacteria were modeled as particles with active Brownian motion, based on and modified from the model developed by Volpe et al. [34]. The main modification was that the finite size of the bacteria was considered when the bacteria got reflected by the surface of the microspheres (Fig. 8A). The microspheres were randomly distributed in the field of view.

**Figure 8:**
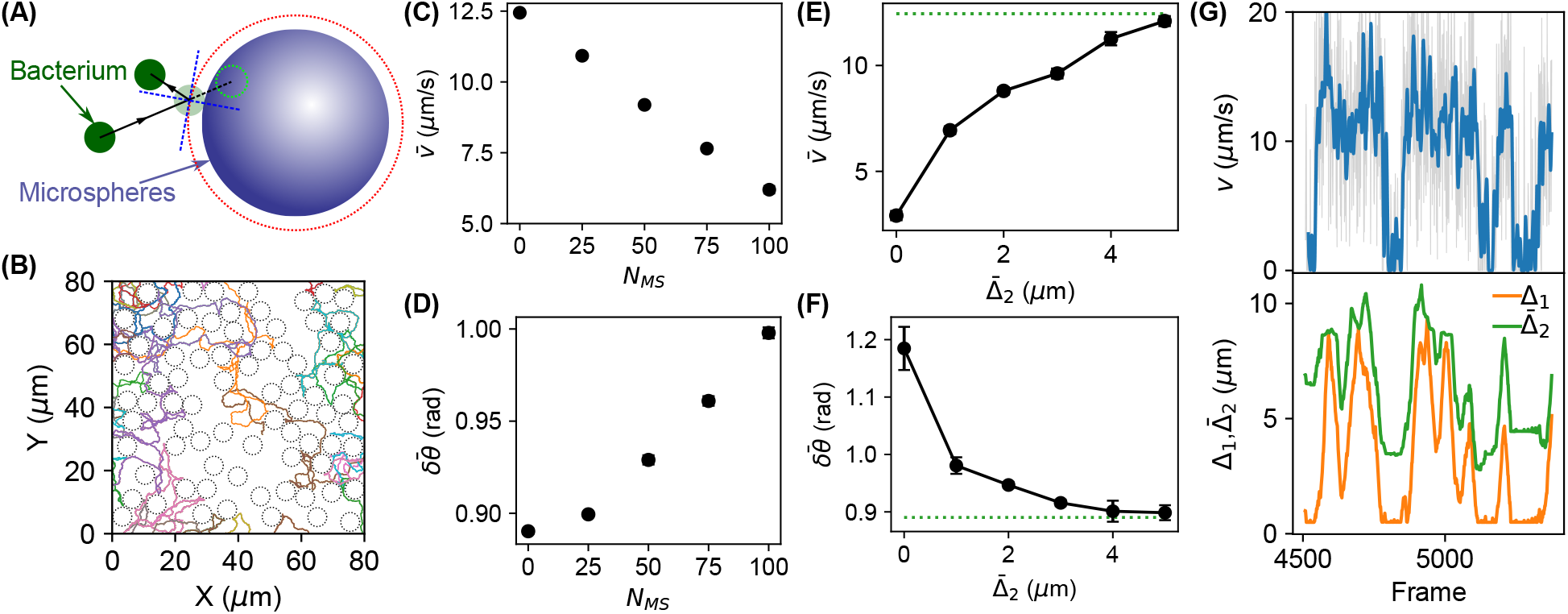
Simulated bacterial motion in the presence of microspheres using the active Brownian motion model. (**A**) The active Brownian motion model while considering the finite size of bacteria and microspheres. (**B**) Examples of simulated bacterial trajectories in the presence of microspheres. (**C**–**D**) Dependence of the (**C**) average bacterial velocity 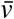 and (**D**) average directional changes of bacteria 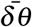 on the number of microspheres *N_MS_*. (**E**–**F**) Dependence of the (**E**) average bacterial velocity 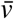 and (**F**) average directional changes of bacteria 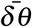 on the average gap size 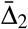. (**G**) An example of the instantaneous velocity (v) of a bacterium and the gap sizes (represented by 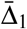 and 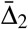), as functions of frame number (time).

Examples of trajectories of simulated bacteria were shown in Fig. 8B, where the simulated bacteria were observed to squeeze through the gaps and pores between the microspheres (Fig. 8B). Similar to the analysis performed for the experimental data, we estimated the instantaneous velocities and directional changes of the bacteria, as well as the corresponding geometric confinements. We found that the simulated results corroborated well with the experimental data. For example, the simulations showed that the average bacterial velocity decreased as the number / density of microspheres increased (Fig. 8C), while the directional change of bacteria increased (Fig. 8D). In addition, as shown in Fig. 8E, the bacterial velocity increased as the geometric confinement became weaker (i.e., as 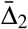 increased), reaching a plateau that corresponded to the average bacterial velocity in the absence of microspheres. The directional change decreased to a plateau as the geometric confinement was weaker (Fig. 8F). Furthermore, we observed bacteria hopping and trapping in the simulations, as well as the positive correlation between the instantaneous bacterial velocity and the geometric confinement (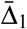 and 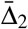), as shown in Fig. 8G. The consistency between the simulated results (Fig. 8) and the experimental observations (Fig. 4–7) suggested that the active Brownian motion model and geometric confinement were sufficient to understand the bacterial motion in microsphere-mimicked porous media.

## Conclusions and Discussions

To summarize, we mimicked porous media, such as soil and tissue, by randomly distributed microspheres at different densities, and investigated the motion of *E. coli* bacteria in such porous media. We observed that the bacterial average velocities were reduced significantly as the density of microspheres increased. In addition, larger directional changes were observed in the presence of microspheres, which mostly happened around the microspheres and due to the collisions with the microspheres. More importantly, the capability of simultaneously localizing both the bacteria and the microspheres allowed us to quantify the distance of the bacteria to the microspheres and the “pore sizes”, and to determine how the observed changes in the bacterial motion depend on the “pore sizes”. It also enabled us to examine individual trajectories and to establish the correlation between the previously observed bacterial hopping trapping in porous media and the geometric confinement imposed by the porous media. Lastly, we ran numerical simulations for active Brownian swimmers in the presence of microspheres and compared the simulated bacterial motion with the experimental results. We observed that the simulated active Brownian swimmers showed lower velocities and higher directional changes as the density of microspheres increased (or the pore sizes decreased), consistent with the experimental data. The numercial simulations also showed positive correlation between bacterial velocity and geometric confinement due to microspheres, consistent with our experimental observations.

Bacterial hopping and trapping were observed and reported previously in 3D porous media made of transparent hydrogel [2, 3]. In the current study, we confirmed similar behaviors of bacteria in quasi-2D porous media formed by microspheres. More importantly, by simultaneously localizing both the microspheres and the bacteria, we established the correlation between bacterial trapping and pore-scale confinement for the first time. We observed that the geometric confinement imposed by microspheres correlated positively with the reduction of bacterial velocity, and negatively with the enhancement of directional changes of bacteria.

It was suggested that bacterial motility fundamentally changes in porous media [2, 3], while other studies implied that the unchanged underlying motility of active Brownian particles predicted the observed bacterial behaviors in heterogenous environments [8–10]. We propose to resolve this controversy by distinguishing the motility and mobility of bacteria (and, in general, active swimmers). Previous experimental results and the current work clearly showed that the movement of bacteria were hindered by the geometric, pore-scale confinements [2, 3, 35]; thus, the ability of bacteria to move easily – bacterial mobility – in porous media was significantly affected by the confinements. However, our study suggested that the active Brownian motion model is sufficient for understanding the experimentall observed motion of bacteria in the absence and presence of microspheres, indicating that the underlying motility – the ability of bacteria to move independently – remains the same in the absence or presence of the microspheres. Therefore, we concluded that the geometric confinment in porous media does not change bacterial motility. It is important to distinguish bacterial motility and bacterial mobility in porous media.

## Acknowledgment

This work was supported by the National Science Foundation (Grant No. 1826642), the National Institute of Food and Agriculture / United States Department of Agriculture (Grant No. 2021-67019-33683), and the Arkansas Biosciences Institute (Grants No. ABI-0189, ABI-0226, ABI-0277, ABI-0326, ABI-2021, and ABI-2022). We are also grateful for support from the Arkansas High Performance Computing Center (AHPCC), which is funded in part by the National Science Foundation (Grants No. 0722625, 0959124, 0963249, and 0918970) and the Arkansas Science and Technology Authority.

## Author Contributions

YW conceived and designed the project; DS performed experiments and acquired the data; AR and AJ contributed to data acquisition and troubleshooting; DS, JO, and YW performed data analysis and visualization; JO and YW performed simulations; DS and YW wrote the paper; all authors reviewed, commented, and revised the paper.

